# Heat inactivation of foetal bovine serum causes protein contamination of extracellular vesicles

**DOI:** 10.1101/2023.03.01.530627

**Authors:** Ornella Urzì, Marta Moschetti, Cecilia Lässer, Junko Johansson, Daniele D’Arrigo, Roger Olofsson Bagge, Rossella Crescitelli

## Abstract

Both the release of extracellular vesicles (EVs) in cell cultures and the cargo that these EVs carry can be influenced by cell culture conditions such as the presence of foetal bovine serum (FBS). Although several studies have evaluated the effect of removing FBS-derived EVs by ultracentrifugation (UC), less is known about the influence of FBS heat inactivation on the cell-derived EVs. To assess this, three protocols based on different combinations of EV depletion by UC and heat inactivation were evaluated, including FBS that was ultracentrifuged but not heat-inactivated, FBS that was heat inactivated before EV depletion, and FBS that was heat inactivated after EV depletion. The FBS samples were then added to the culture media of three melanoma cell lines, and after 72 h both large and small EVs were isolated by differential UC. We demonstrated by transmission electron microscopy, protein measurement, and quantification of the number of particles that heat inactivation performed after EV depletion reduced the purity of small EVs but had no effect on large EV purity. Quantitative mass spectrometry analysis of FBS-derived small EVs showed that the EV protein content was different when FBS was heat inactivated after EV depletion compared to EVs isolated from FBS that was not heat inactivated or that was heat inactivated prior to EV depletion. Moreover, several of the quantified proteins were wrongly attributed to be of human origin because the EVs were of obvious bovine origin. Our results demonstrated that proteins of bovine origin coming from FBS-derived EVs could mistakenly be attributed to human cell-derived EVs in EV proteomic studies. Moreover, we concluded that heat inactivation performed after EV depletion induced the release of proteins that might contaminate EV samples, and the recommendation is therefore to always perform heat inactivation prior to EV depletion.

## 1 INTRODUCTION

Extracellular vesicles (EVs) are lipid-enclosed particles released by all cell types into the extracellular space^1^. Although EVs were initially considered garbage bags^2, 3^, they now represent one of the most studied mechanisms of cell–cell communication^4^. EVs can be found in several biological fluids, including, blood, urine, saliva, and cerebrospinal fluid^5–8^. Based on their size or biogenesis, several subtypes of EVs have been identified such as exosomes, microvesicles, and apoptotic bodies^9, 10^. The existing overlap between different EV subpopulations is still debated by the EV community, but EVs are commonly divided into two major classes based on size, namely small EVs (sEVs, 30–200 nm) and large EVs (lEVs, 200–1000 nm)^11–13^. The cargo of EVs consists of proteins, nucleic acids (e.g. DNA and both coding and non-coding RNA), and lipids, and these vary depending on the releasing cell^14^. The bioactive molecules carried by EVs can enter target cells and modify their phenotype and metabolism^4^. The EVs derived from cell lines are the most-studied source of EVs^15, 16^. However, the culture conditions and especially the use of foetal bovine serum (FBS) may influence the results of using EVs obtained in *in vitro* studies^17^.

FBS is widely used to supplement culture media because it provides growth factors, amino acids, and vitamins and promotes cell growth and adhesion as well as it protects cells from stressful conditions^18^. Although most cells need FBS to grow properly in culture, the use of FBS carries some disadvantages, mainly related to undefined composition^19, 20^ and ethical concerns^21, 22^. Moreover, because FBS can contain microbial contaminants such as mycoplasma, viruses, and prion proteins^22^ it is usually heat-inactivated at 56°C for 30–60 min before use^23, 24^. The heat inactivation also inactivates complement activity and thus is a widely used practice, especially for some cell types, such as immune cells, that can be affected by complement activation^23, 25^.

It has been widely demonstrated that FBS contains large amounts of EVs that might influence the results of EV analysis in *in vitro* studies^26^. Bovine EVs represent an important limit in the use of FBS as a supplement for cell culture media because they can contaminate cell-derived EV samples^27^ and can influence the growth and phenotype of cultured cells^28, 29^. For these reasons, when performing EV studies it is recommended to deplete the EVs from the FBS^17, 30, 31^. The most commonly used protocol to deplete EVs from FBS is ultracentrifugation (UC), which consists of centrifuging the FBS at 100,000–120,000 × *g* for 1–18 h and then discarding the pellet of bovine-derived EVs^26, 28, 30^.

Although UC protocols have been optimized in recent years, some studies suggest that UC does not completely eliminate all FBS-derived EVs^28, 32, 33^. This issue has led to the development of new protocols for EV depletion such as ultrafiltration^34^ and polymer-based precipitation^35^, which seem promising although further studies are needed to improve them.

An aspect that has been neglected in the optimization and evaluation of EV-depletion protocols is the heat inactivation procedure. Generally, authors do not mention if the FBS was heat inactivated in their studies^36, 37^, and even when this is mentioned the time and temperature of the procedure are often missing^32, 38^. Moreover, only rarely do authors specify whether the heat inactivation was performed before or after EV depletion by UC. Because FBS is known to potentially contaminate cell-derived EV samples, the aim of this study was to determine the effect of heat inactivation on EV purity using quantitative mass spectrometry.

## 2 METHODS

### 2.1 Preparation of EV-depleted FBS and FBS heat inactivation

The FBS purchased from Sigma-Aldrich (St louis, MO, lot number: 0001638271) was cleaned of EVs following three slightly different protocols based on different combinations of UC and heat inactivation. The UC step was an EV depletion step where the FBS was ultracentrifuged at 118,000 × *g* (Type 45 Ti rotor, 38,800 rpm, Beckman Coulter, Brea, CA) for 18 h at 4°C followed by a filtration step (0.22 μm). In the heat inactivation step the FBS was placed in a water bath at 56°C for 30 min under agitation. Three different combinations of UC and heat inactivation were tested and are described in Fig. 1, and the protocol was the same for all conditions except for the time when the heat inactivation was performed: (a) FBS was ultracentrifuged but not heat-inactivated at all (no-HI FBS), (b) FBS was heat-inactivated prior to EV depletion (HI-before EV-depl FBS), and (c) FBS was heat inactivated after EV depletion (HI-after EV-depl FBS). The supernatant was filtered through a 0.22 μm filter and then stored at −20°C until use.

**FIGURE 1.**
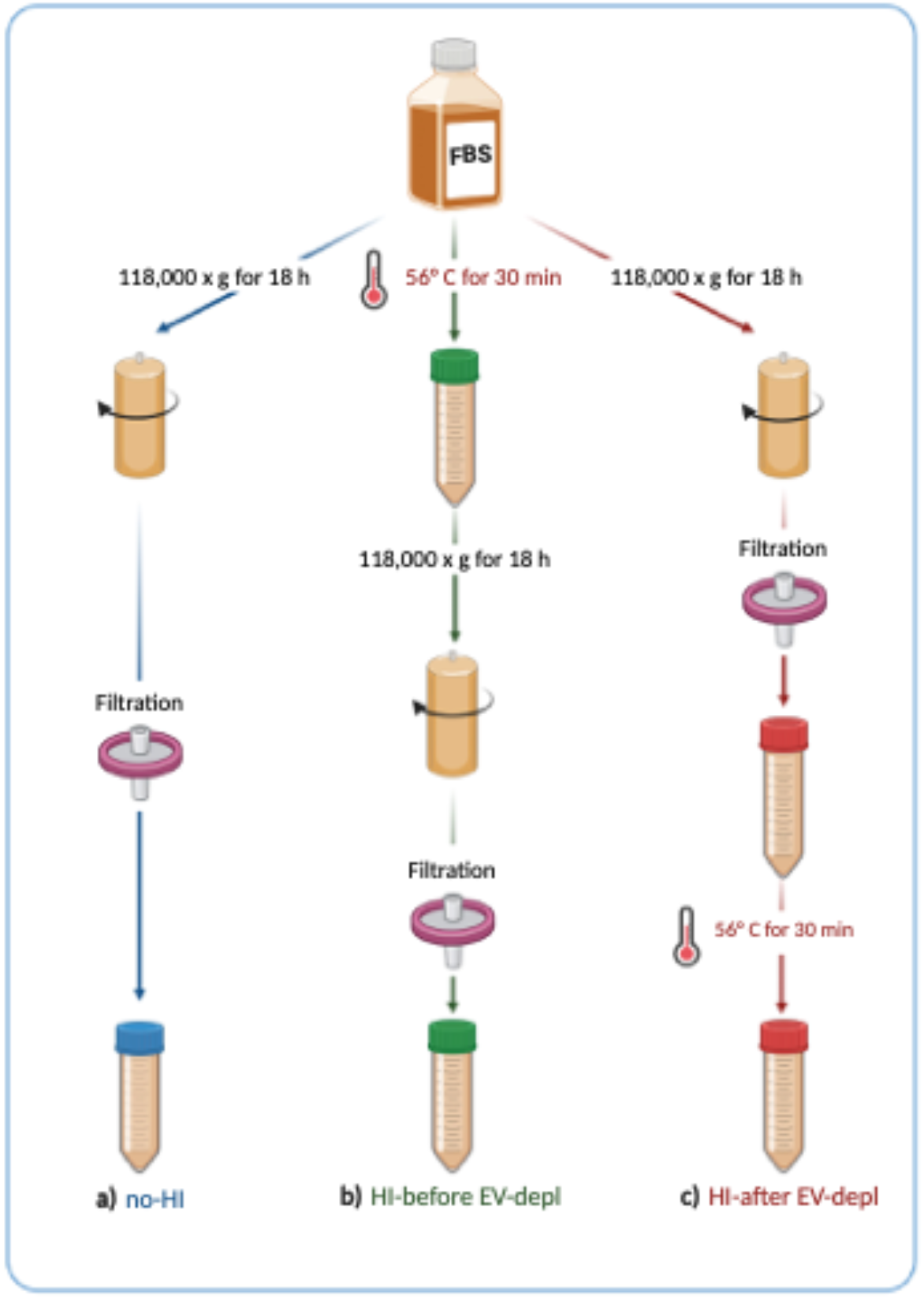
Preparation of EV-depleted FBS with three different combinations of UC and heat inactivation. Created with BioRender.com

### 2.2 Cell cultures

The human skin melanoma cell line MML-1 (CLS, Eppelheim, Germany) and the human uveal melanoma cell lines UM22Bap1^+/+^ and UM22Bap1^-/-^ were used for the isolation of sEVs and 1EVs. UM22Bap1^+/+^ and UM22Bap1^-/-^ are cell lines isolated from the liver of a patient affected by uveal melanoma with liver metastasis and a homozygous frameshift deletion in *BAP1*. UM22Bap1^+/+^ cells were transfected using a retroviral vector with a functional copy of *BAP1*, while UM22Bap1^-/-^ cells received an empty control vector^39^. MML-1, UM22Bap1^+/+^, and UM22Bap1^-/-^ cells were cultured in RPMI-1640 (Cytiva, Loga, UT, USA) supplemented with 10% FBS (Sigma-Aldrich, St louis, MO), 100 units/ml penicillin, 100 mg/ml streptomycin (Cytiva), and 2 mM L-glutamine (Cytiva). The cells were grown in three different conditions: (a) media supplemented with no-HI FBS, (b) media supplemented with HI-before EV-depl FBS, and (c) media supplemented with HI-after EV-depl FBS. Cells were seeded at 50,000 cells/cm^2^ and grown in a 37°C humidified incubator with 5% CO_2_. Cells were passaged every third day, and cell viability was assessed using the trypan blue exclusion method.

### 2.3 Cell viability assays

In order to assess the viability of the cells grown in the three described conditions, cells from three melanoma cell lines (MML-1, UM22Bap1^+/+^, and UM22Bap1^-/-^) were seeded in flat-bottomed 6-well plates and incubated at 37°C with 5% CO2 for 72 h. The cell number was determined manually using a trypan blue exclusion assay and an optical microscope, and 5 × 10^5^ cells/well were seeded in complete medium containing FBS processed by one of the three protocols. After 72 h the cells were photographed with an Olympus CKX41 optical microscope equipped with an Infinity 1 camera (Luminera, Ottawa, ON) using the Infinity Analyze software (version 6.5.0, Luminera). The cells were gently detached from the cell culture plate with 0.25% trypsin at 37°C for approximately 3 min. After washing, one aliquot of each cell line was used for manual counting of the cell number, while the rest of the cells were stained with a Live/Dead Fixable Far Red Dead Cell Stain Kit (Invitrogen, Oregon, US) in phosphate-buffered saline (PBS) and analysed on a BD LSRFortessa instrument (BD Biosciences, San Jose, CA). Subsequent data analysis was performed in BD FACSDiva Software version 9.0.1 (BD Biosciences).

### 2.4 Isolation of EVs

Both lEVs and sEVs were isolated from the conditioned media of MML-1, UM22Bap1 ^+/+^, and UM22Bap1^-/-^ cell lines as described previously^11^. Briefly, the conditioned media was collected after 72 h and subjected to differential centrifugations. Cells and debris were removed by centrifugations at 300 × *g* for 10 min and at 2,000 × *g* for 20 min at 4°C. The supernatant was centrifuged at 16,500 × *g* for 20 min at 4°C and 118,000 × *g* for 2.5 h at 4°C to collect lEVs and sEVs, respectively (Type 45 Ti rotor (Beckman Coulter) at 14,500 rpm and 38,800 rpm and with a k-factor of 1279.1 and 178.6, respectively). Pellets were resuspended in PBS and stored at –80°C for further experiments. The same EV isolation protocol was performed on FBS after the EV depletion and heat inactivation as described in Fig. S4.

### 2.5 Protein measurement

The protein concentrations of the lEVs and sEVs isolated from cell cultures and FBS were measured by Qubit (Thermo Fisher Scientific) according to the manufacturer’s protocol.

### 2.6 Nanoparticle tracking analysis

The numbers of particles were measured using ZetaView PMX110 (Particle Metrix, Meerbusch, Germany) as described previously^11^. Measurements were done at all 11 positions, and the camera sensitivity was set to 80 and the shutter was set to 100. The chamber temperature was automatically measured and integrated into the calculation. Data were analysed using the ZetaView analysis software version 8.2.30.1 with a minimum and maximum size of 5 nm and 5000 nm, respectively, and minimum brightness of 20. Three biological replicates were measured from each sample.

### 2.7 Transmission electron microscopy (TEM)

Investigation of EVs by negative staining was performed as previously described^40, 41^. Briefly, 5 μg of EVs was placed onto glow discharged 200-mesh formvar/carbon copper grids (Electron Microscopy Sciences, Hatfield Township, PA). The EVs were washed two times with water and then fixed in 2.5% glutaraldehyde. After two further washes in water, the samples were stained with 2% uranyl acetate for 1.5 min. Negative-stained samples were examined on a Talos L120C electron microscope (Thermo Fisher Scientific) at 120 kV with a CCD camera.

### 2.8 Sample preparation and digestion for mass spectrometry

The proteomic analysis was performed at The Proteomics Core Facility at Sahlgrenska Academy, Gothenburg University. The samples and reference pools (representative reference material containing equal amounts of all the samples) were digested using a modified filter-aided sample preparation method^42^. In short, samples (35 μg) were reduced with 100 mM dithiothreitol at 60°C for 30 min, transferred to Microcon-30kDa Centrifugal Filter Units (Merck), and washed several times with 8 M urea and once with digestion buffer (50 mM TEAB, 0.5% sodium deoxycholate) prior to alkylation with 10 mM methyl methanethiosulfonate in digestion buffer for 30 min at room temperature. Samples were digested with trypsin (Pierce MS grade Trypsin, Thermo Fisher Scientific, ratio 1:100) at 37°C overnight, and an additional portion of trypsin was added and incubated for another 2 h. Peptides were collected by centrifugation and labelled using TMT 10-plex isobaric mass tagging reagents (Thermo Fisher Scientific) according to the manufacturer’s instructions. The samples were combined into one TMT-set, and sodium deoxycholate was removed by acidification with 10% TFA. The TMT-set was further purified using High Protein and Peptide Recovery Detergent Removal spin columns and Pierce peptide desalting spin columns (both from Thermo Fischer Scientific) according to the manufacturer’s instructions prior to basic reversed-phase chromatography (bRP-LC) fractionation. Peptide separation was performed using a Dionex Ultimate 3000 UPLC system (Thermo Fischer Scientific) and a reverse-phase XBridge BEH C18 column (3.5 μm, 3.0 mm × 150 mm, Waters Corporation) with a gradient from 3% to 100% acetonitrile in 10 mM ammonium formate at pH 10.00 over 23 min at a flow rate of 400 μL/min. The 40 fractions were concatenated into 20 fractions, dried, and reconstituted in 3% acetonitrile and 0.2% formic acid.

### 2.9 nanoLC-MS/MS analysis and database searching

Each fraction was analysed on an Orbitrap Lumos Tribrid mass spectrometer equipped with the FAIMS Pro ion mobility system interfaced with an nLC 1200 liquid chromatography system (all from Thermo Fisher Scientific). Peptides were trapped on an Acclaim Pepmap 100 C18 trap column (100 μm × 2 cm, particle size 5 μm, Thermo Fischer Scientific) and separated on an in-house-constructed analytical column (350 mm × 0.075 mm I.D.) packed with 3 μm Reprosil-Pur C18-AQ particles (Dr. Maisch, Germany) using a gradient from 3% to 80% acetonitrile in 0.2% formic acid over 85 min at a flow rate of 300 nL/min. The FAIMS Pro alternated between the compensation voltages of –40 V and –60 V, and essentially the same data-dependent settings were used at both compensation voltages. Precursor ion mass spectra were acquired at 120,000 resolution, a scan range of 450–1375, and a maximum injection time of 50 ms. MS2 analysis was performed in a data-dependent mode, where the most intense doubly or multiply charged precursors were isolated in the quadrupole with a 0.7 m/z isolation window and dynamic exclusion within 10 ppm for 60 s. The isolated precursors were fragmented by collision-induced dissociation at 35% collision energy with a maximum injection time of 50 ms for 3 s (‘top speed’ setting) and detected in the ion trap. This was followed by multinotch (simultaneous) isolation of the top-10 MS2 fragment ions within the m/z range 400–1200, fragmentation (MS3) by higher-energy collision dissociation at 65% collision energy, and detection in the Orbitrap at 50,000 resolution in the m/z range 100–500 and a maximum injection time of 105 ms.

The data files for each set were merged for identification and relative quantification using Proteome Discoverer version 2.4 (Thermo Fisher Scientific). The search was against *Homo sapiens* (Swissprot Database March 2022) and Bovine (Swissprot Database April 2021) using Mascot 2.5 (Matrix Science) as the search engine with a precursor mass tolerance of 5 ppm and a fragment mass tolerance of 0.6 Da. Tryptic peptides were accepted with one missed cleavage, variable modifications of methionine oxidation, and fixed cysteine alkylation, and TMT-labelled modifications of N-termini and lysines were selected. Percolator was used for PSM (peptide spectral match) validation with the strict FDR threshold of 1%. TMT reporter ions were identified with a 3 mmu mass tolerance in the MS3 higher-energy collision dissociation spectra, and the TMT reporter abundance values for each sample were normalized to the total peptide amount. Only the quantitative results for the unique peptide sequences with a minimum SPS match of 50% and an average S/N above 10 were taken into account for the protein quantification. A reference sample was used as the denominator for calculating the ratios. The quantified proteins were filtered at 1% FDR and grouped by sharing the same sequences in order to minimize redundancy.

### 2.10 Bioinformatics and statistical analysis

Where appropriate, data are expressed as the mean and standard deviation of the mean (SD). Statistical analysis was performed by non-paired Student’s t-test or one-way ANOVA for multiple comparisons in GraphPad Prism 6 (GraphPad Software Inc., La Jolla, CA).

Qlucore Omics Explorer (Qlucore, Lund, Sweden) was used for the principal component analysis. The proteins that were enriched in the FBS HI-after EV-depl were analysed using the Database for Annotation, Visualization and Integrated Discovery (DAVID; http://david.abcc.ncifcrf.gov/ [accessed: 19–07–2022]) to determine the cellular components of the proteins.

## 3 RESULTS

### 3.1 FBS heat inactivation causes contamination of sEVs isolated from cell cultures

To compare the EV yield and purity, we cultured three melanoma cell lines, MML-1, UM22Bap1 ^+/+^, and UM22Bap1^-/-^, in media supplemented with no-HI FBS, HI-before EV-depl FBS, or HI-after EV-depl FBS. Because several cell culture parameters can affect EV release^17^, we first confirmed that cell morphology and confluence were not altered in the three culture conditions (Fig. S1). Moreover, cell viability was evaluated at seeding time (time 0) and after 72 h (when the cell supernatant was harvested to collect EVs) using trypan blue, and the number of live cells was analysed by flow cytometry. The cell viability was similar in all of the experimental conditions for the three cell lines, suggesting that the different combinations of EV depletion and heat inactivation did not affect cell viability (Fig. 2a-c). Similarly, the doubling time was comparable in media supplemented with no-HI FBS, HI-before EV depl FBS, and HI-after EV depl FBS in all three cell lines (Fig. 2d-f). Overall, these results demonstrated that media supplemented with no-HI FBS, HI-before EV-depl FBS, and HI-after EV-depl FBS did not affect the morphology or growth of the MML-1, UM22Bap1^+/+^, and UM22Bap1^-/-^ cell lines.

**FIGURE 2.**
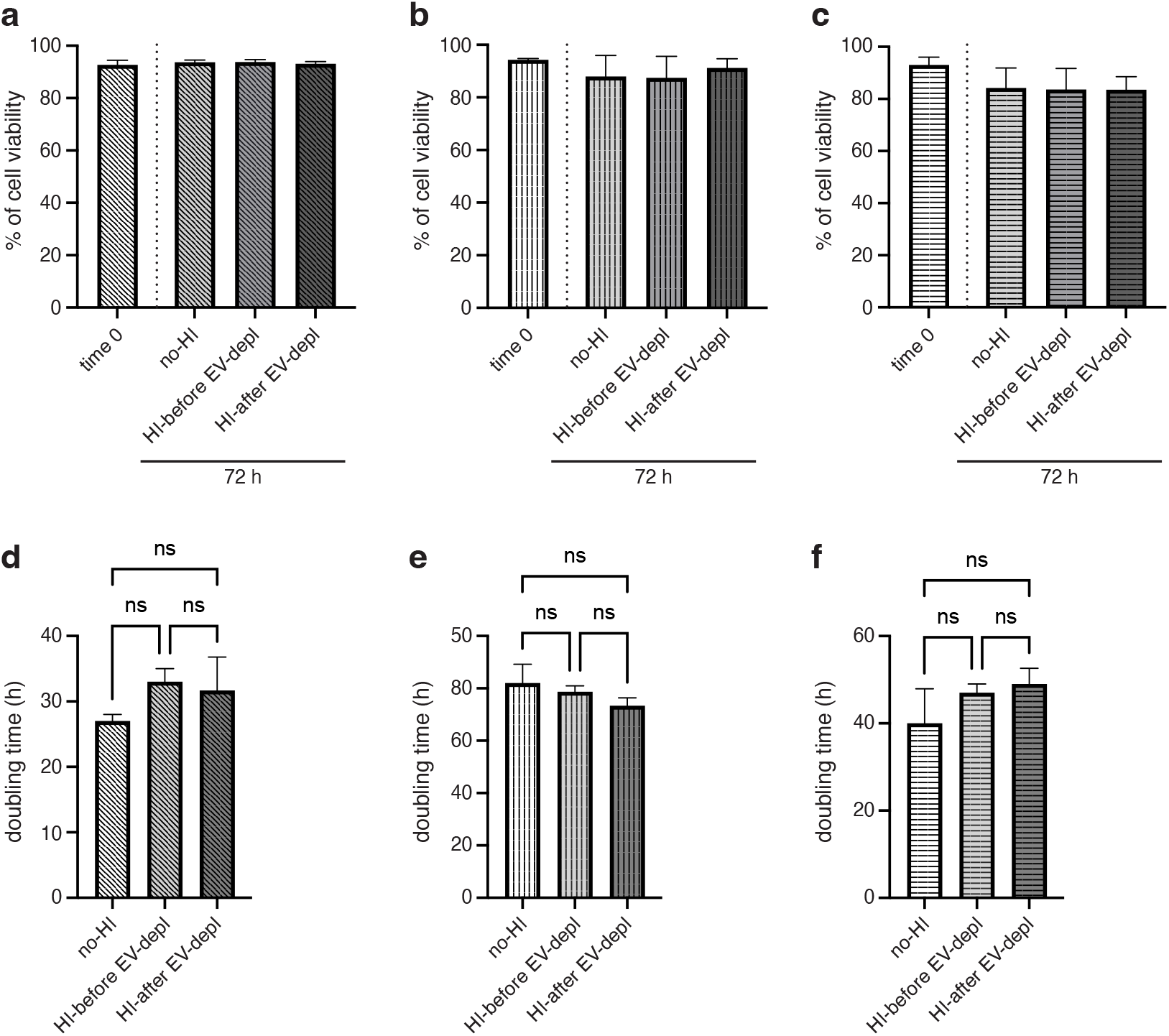
Analysis of cell viability. Percentage of cell viability assessed by cell counting using trypan blue of (a) MML-1, (b) UM22Bap1^+/+^, and (c) UM22Bap1^-/-^ cells in media supplemented with no-HI FBS, HI-before EV-depl FBS, and HI-after EV-depl FBS at time 0 and after 72 h. The doubling time (h) of (d) MML-1, (e) UM22Bap1^+/+^, and (f) UM22Bap1^-/-^ cells in media supplemented with no-HI FBS, HI-before EV-depl FBS, and HI-after EV-depl FBS. *N* = 3. The results are presented as the average ± SD. Ns = non-significant.

We next isolated and characterized EVs released by the three different cell lines using the three different conditions. The three melanoma cell lines were seeded in media supplemented with no-HI FBS, HI-before EV-depl FBS, and HI-after EV-depl FBS, and after 72 h the conditioned media were collected. Both lEVs and sEVs were isolated by differential centrifugations and the EVs were analysed. TEM analysis demonstrated that both lEVs and sEVs showed typical morphology and a clear difference in size (mean diameter 120 nm vs. 80 nm, respectively) in all conditions analysed (MML-1 in Fig. 3a and UM22Bap1^+/+^ and UM22Bap1^-/-^ in Fig. S2). However, elements surrounding the sEVs (a strong image background) were visible in picture of sEVs isolated from MML-1 cells grown in media supplemented with HI-after EV-depl FBS but not in image of sEVs isolated from cells grown in media supplemented with no-HI FBS and HI-before EV-depl FBS (Fig. 3a). Similar results were obtained for sEVs derived from UM22Bap1^+/+^ and UM22Bap1^-/-^ cells (Fig. S2). Moreover, the protein measurement showed an increased amount of protein in sEVs isolated from the media supplemented with HI-after EV-depl FBS compared to the sEVs isolated from media containing no-HI FBS and HI-before EV-depl FBS. This result was different in lEVs in which no differences in protein levels were observed (Fig. S3 a-c). The evaluation of EV purity, calculated as the numbers of particles per μg protein, showed that sEVs isolated from cells grown in HI-after EV-depl FBS had lower purity compared to sEVs isolated from cells grown in no-HI FBS and HI-before EV depl FBS. No significative effect on the purity of lEVs was observed. Similar results were obtained from EVs isolated from all three of the cell lines that were investigated (Fig. 3b-c). Together these results indicated that heat inactivation of FBS induces the production of contaminants that could affect the purity of cell-derived sEV samples if the heat inactivation is performed after EV depletion by UC.

**FIGURE 3.**
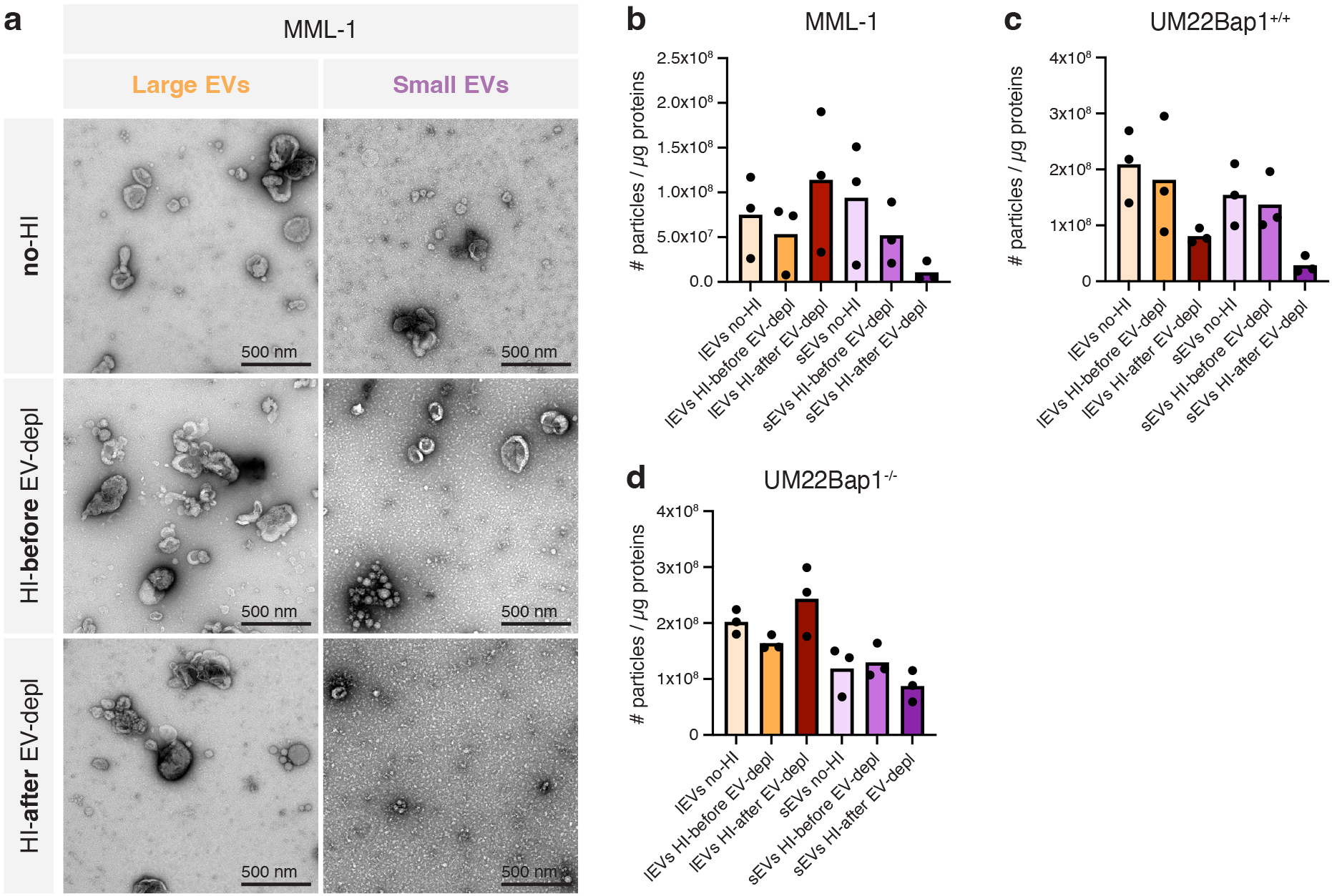
Analysis of EVs isolated from melanoma cells cultured in media supplemented with no-HI FBS, HI-before EV-depl FBS, and HI-after EV-depl FBS. (a) TEM analysis of EVs isolated from MML-1 cells cultured in media supplemented with no-HI FBS, HI-before EV-depl FBS, and HI-after EV-depl FBS. Five micrograms of lEVs and sEVs were loaded onto grids, negative stained, and evaluated by TEM. Scale bars = 500 nm. The ratio of the particle number to the protein concentration of lEVs and sEVs isolated from (b) MML-1, (c) UM22Bap1^+/+^, and (d) UM22Bap1^-/-^ cells in media supplemented with no-HI FBS, HI-before EV-depl FBS, and HI-after EV-depl FBS. The results are presented as the mean, but individual values from three different experiments are shown as well. *N* = 3.

### 3.2 FBS heat inactivation is the cause of sEV sample contamination

Given that we did not observe any changes in cell phenotype or viability and that the only difference between the three experimental conditions was how the FBS was handled prior to being added to the culture media, we next determined what the FBS contributed to the EV samples isolated from the cell cultures. The three types of pre-processed FBS were centrifuged to isolate FBS-derived lEVs and sEVs as described in Fig S4. The sEV pellet was easily visualized when isolated from HI-after EV-depl FBS, but this was not the case for the sEV and lEV pellets isolated from no-HI or HI-before EV-depl FBS (Fig. 4a, left panel). TEM analysis of FBS-derived EV pellets (both lEVs and sEVs) showed that 18 h of UC did not completely eliminate bovine EVs, confirming what was demonstrated previously^28, 32^. Moreover, a strong background due to elements surrounding the EVs was visible in TEM images of sEVs isolated from HI-after EV-depl FBS, likely due to protein contaminants. The strong background was not present in images of lEVs and sEVs isolated from no-HI FBS or HI-before EV-depl FBS (Fig. 4a, right panel). The background observed in sEV TEM images was likely due to protein contaminants as demonstrated by the greater amount of protein in this samples compared to the other samples (Fig. S5).

**FIGURE 4.**
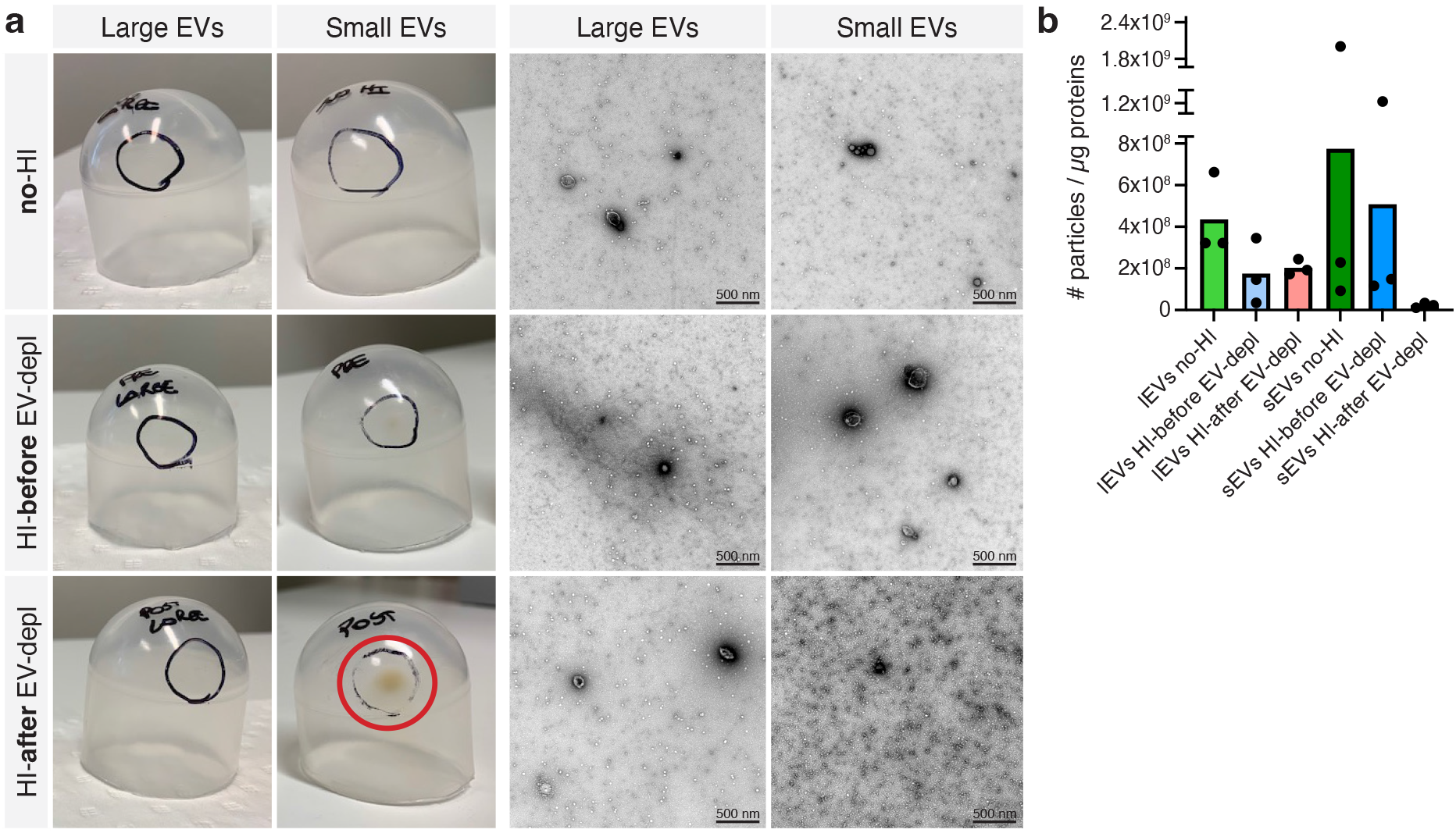
Analysis of lEVs and sEVs isolated by the differential centrifugation of no-HI FBS, HI-before EV-depl FBS, and HI-after EV-depl FBS. (a, left panel) Representative pictures of pellets obtained by the UC of no-HI FBS, HI-before EV-depl FBS, and HI-after EV-depl FBS. An obviously visible pellet isolated from HI-after EV-depl FBS is indicated (red circle). (a, right panel) TEM images of lEVs and sEVs isolated from no-HI FBS, HI-before EV-depl FBS, and HI-after EV-depl FBS. (b) The ratio of the particle number and protein concentration of lEVs and sEVs isolated from no-HI FBS, HI-before EV-depl FBS, and HI-after EV-depl FBS. The results are presented as the mean, but individual values from three different experiments are shown. *N* = 3.

Similar to cell-derived sEVs, the number of particles per μg of protein in sEVs isolated from HI-after EV-depl FBS was lower compared to no-HI FBS and HI-before EV-depl FBS indicating the reduced purity of this pellet compared to the other pellets (Fig. 4b). To further confirm that FBS was the source of sEV contamination, we also performed EV isolation from media supplemented with no-HI FBS, HI-before EV-depl FBS, and HI-after EV-depl FBS, but without ever being added to the cells. The results were consistent with previous observations, and sEVs obtained from media supplemented with HI-after EV-depl FBS were enriched in contaminants and their purity was lower compared to sEVs isolated from media supplemented with no-HI FBS and HI-before EV-depl FBS. No EVs were collected from the cellular media not supplemented with FBS (Fig. S6). Overall, these results indicated that heat inactivation of FBS after EV depletion by UC induced the release of contaminants that ended up in the sEV pellet. Consequently, the sEVs isolated from the cell lines were contaminated by sEVs remaining in FBS after 18 h of centrifugation and by contaminants produced during the heat inactivation process if it was performed after EV depletion.

### 3.3 Quantitative proteomic analysis revealed that heat inactivation performed after EV depletion could alter the protein content of cell-derived sEVs

To further investigate the nature of the contaminants induced by the heat inactivation process, we performed quantitative proteomics on sEV pellets isolated from FBS using the three protocols (Fig. S4). In total, we identified 2074 proteins and quantified 1879 proteins (Fig. S7a and b respectively). Among the quantified proteins, 893 were of human origin, 870 were of bovine origin, 91 were of both human and bovine origin, and 25 were not of human or bovine origin (Fig. S7b). Furthermore, we manually compared the 893 human proteins and the 870 bovine proteins and found that 36 proteins were present in both lists (Supplementary Table 1). Because the FBS-derived sEV pellets were obviously of bovine origin, we initially focused our analysis on the bovine protein data set. A principal component analysis including all quantified bovine proteins was performed to visualize the relationship between the sEV pellets isolated under the three different conditions. Component 1, representing the largest variability with 53%, distinguished the HI-after EV-depl FBS-derived sEVs from the no-HI FBS-derived sEVs and HI-before EV-depl FBS-derived sEVs, indicating that the pellet isolated from HI-after EV-depl FBS was different from the pellets isolated from no-HI FBS and HI-before EV depl FBS (Fig. 5a). Proteins quantified in sEVs from no-HI FBS and HI-before EV-depl FBS were similar as shown by volcano plot analysis (Fig. 5b), but were different from the protein quantified in HI-after EV-depl FBS (Fig. 5c, d). These results suggested that heat inactivation may alter the protein content of FBS and that these proteins end up within the sEV pellet during UC, thus leading to an overestimation of the protein measured in sEVs isolated from cultured cells as well as mis-interpretation of the proteomics results.

**FIGURE 5.**
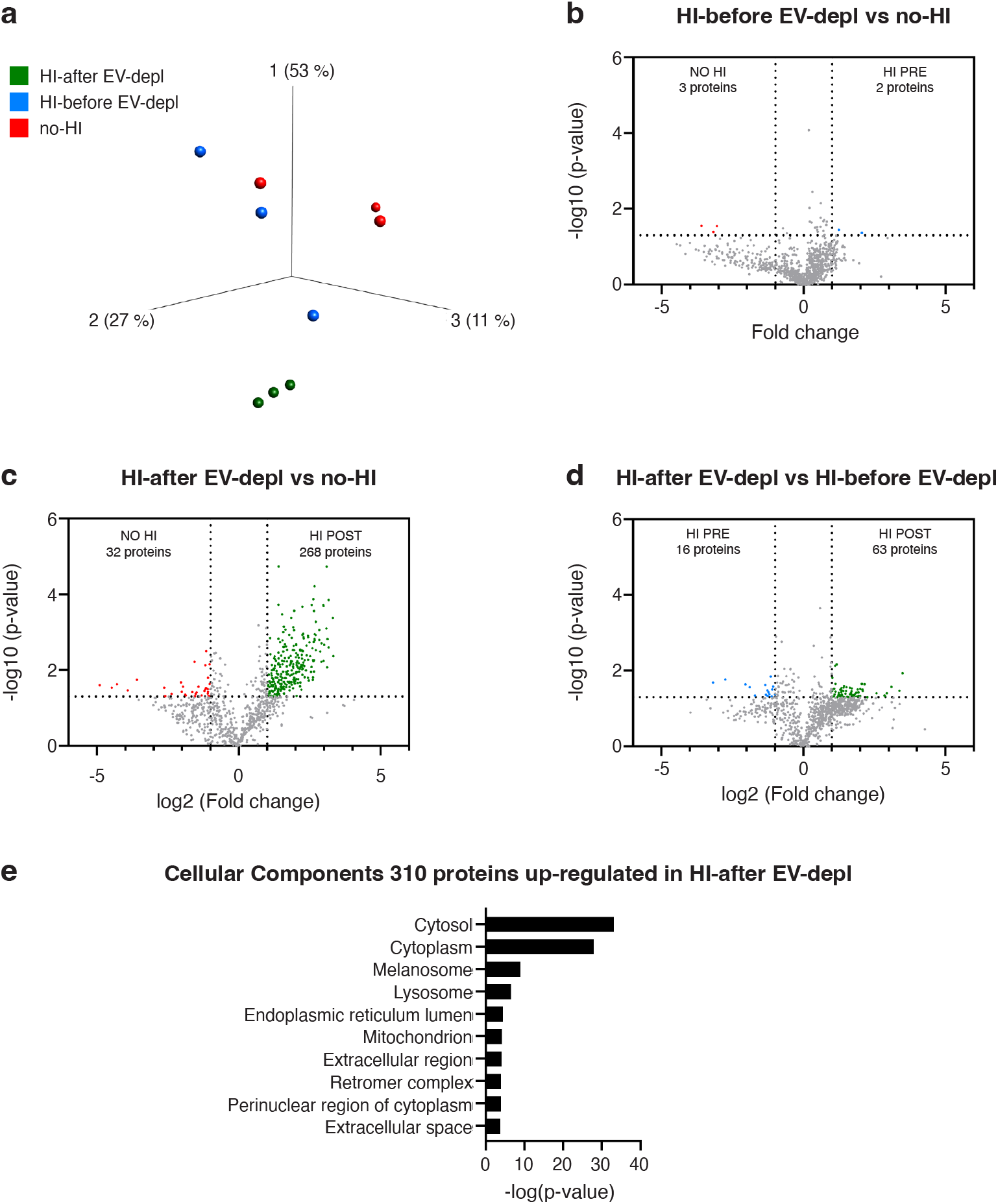
Analysis of sEVs isolated from FBS against the bovine dataset. Quantitative proteomics (TMT) was used to determine the differences in the sEVs isolated from no-HI FBS, HI-before EV-depl FBS, and HI-after EV-depl FBS, *N* = 3. (a) Principal component analysis illustrating the relation between sEVs isolated from no-HI FBS (red), HI-before EV-depl FBS (blue), and HI-after EV-depl FBS (green). (b-d) Volcano plot showing proteins with differential expression between sEVs from no-HI FBS, HI-before EV-depl FBS, and HI-after EV-depl FBS (b), sEVs from HI-after EV-dep FBS and no-HI FBS (c), and sEVs from HI-after EV-dep FBS and HI-before EV-depl FBS (d). Grey dotted lines show p-value < 0.05 and two-fold change cut-offs. (e) Database for Annotation, Visualization and Integrated Discovery (DAVID) was used to determine the most enriched cellular compartments associated with proteins that were significantly enriched in FBS HI-after EV-depl compared to FBS no-HI. The 10 most enriched terms (based on p-value) are displayed (310 proteins, p-value = 0.001, q-value = 0.009).

We then focused on the proteins enriched in the sEV pellet isolated from HI-after EV-depl FBS, and we specifically analysed the top 20 up-regulated proteins in the sEV pellet from HI-after EV-depl FBS versus no-HI FBS (Fig. S7c) and versus HI-before EV-depl FBS (Fig. S7d). Most of these proteins were involved in metabolism, such as beta-enolase (ENO3), cytoplasmic aconitate hydratase (ACO1), and fatty acid synthase (FASN). Interestingly, we also identified exocyst complex component 8 (EXOC8), which belongs to the family of exocytosis complexes and plays a critical role in vesicular trafficking and promotes the fusion of post-Golgi vesicles to the plasma membrane^43^. The gene ontology (GO) terms analysis revealed that sEVs from HI-after EV-depl FBS were enriched in proteins that were associated with the cytosol and lysosomes as well as proteins associated with the extracellular region and the extracellular space (Fig. 5e). These results suggested that heat inactivation performed after EV depletion by UC modified the proteins in the FBS that then copelleted with cell-derived sEVs.

### 3.4 Proteins in EVs from heat inactivated FBS after EV depletion could be mistakenly attributed to human cell-derived EVs

Because the majority of the studies analysing proteomics of EVs have focused on proteins of human origin, and thus have investigated the human protein list, we also analysed the proteomic results against the human database. A principal component analysis was performed including all quantified human proteins (Fig. 6a). Component 1 represented the largest variability (47%) and distinguished sEVs obtained from HI-after EV-depl FBS from those obtained from no-HI FBS and HI-before EV-depl FBS. A strong similarity was observed in proteins of sEVs from no-HI FBS and HI-before EV-depl FBS (Fig. 6b). In contrast, the proteins in sEVs from HI-after EV-depl FBS showed significant differences compared to sEVs from no-HI (Fig. 6c) and HI-after EV-depl FBS (Fig. 6d). These results were similar to the results obtained from the analysis of bovine proteins (Fig. 5 a-d), highlighting that heat inactivation after UC affected the protein content of FBS and that these proteins ended up in the sEV pellet during the EV depletion process by UC thus contaminating the cell-derived sEV samples.

**FIGURE 6.**
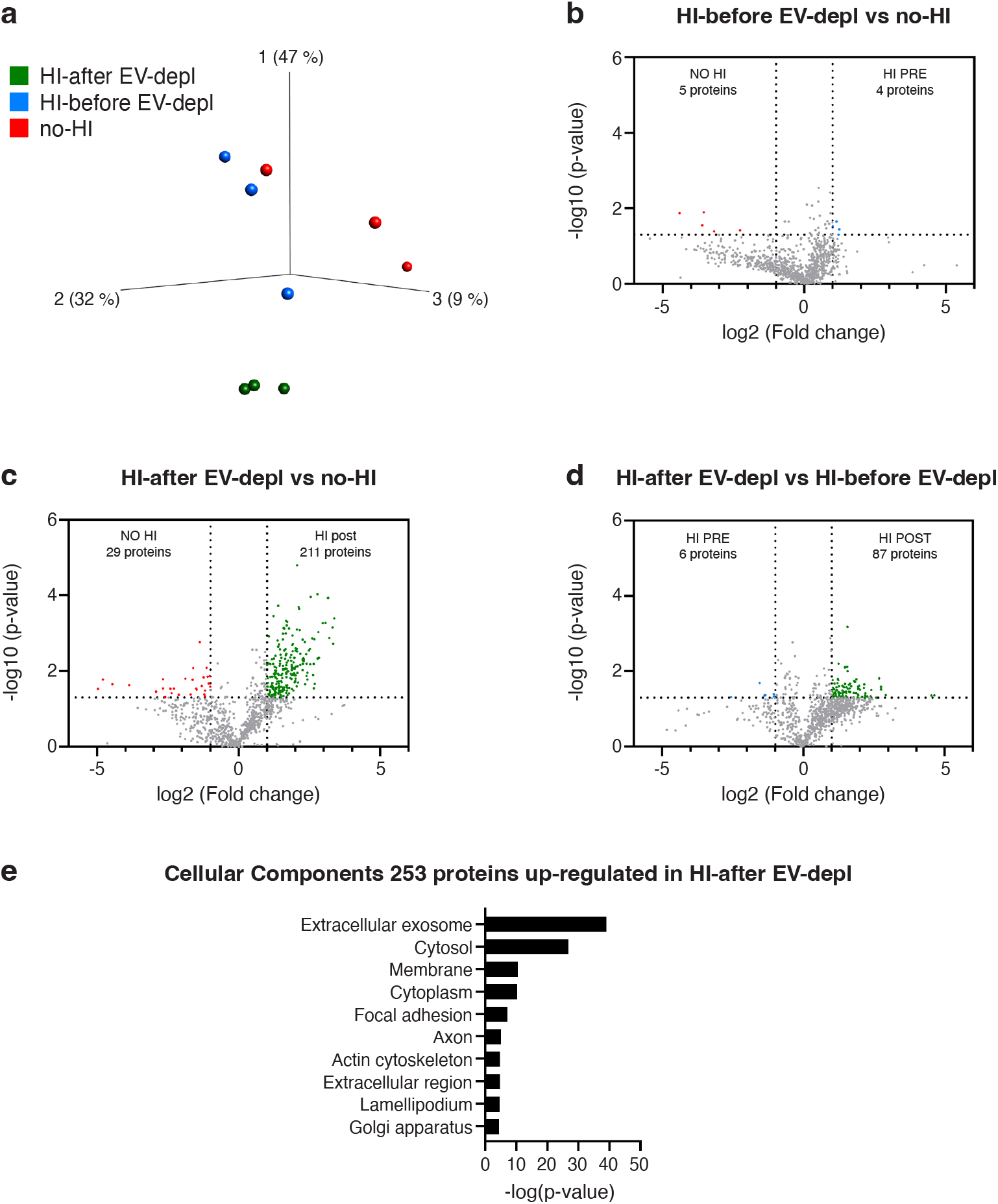
Analysis of sEVs isolated from FBS against the human dataset. Quantitative proteomics (TMT) was used to determine the differences in the sEVs isolated from no-HI FBS, HI-before EV-depl FBS, and HI-after EV-depl FBS, *N* = 3. (a) Principal component analysis illustrating the relation between sEVs isolated from no-HI FBS (red), HI-before EV-depl FBS (blue), and HI-after EV-depl FBS (green). (b-d) Volcano plot showing proteins with differential expression between sEVs from no-HI FBS, HI-before EV-depl FBS, and HI-after EV-depl FBS (b), sEVs from HI-after EV-dep FBS and no-HI FBS (c), and sEVs from HI-after EV-dep FBS and HI-before EV-depl FBS (d). Grey dotted lines show p-value < 0.05 and two-fold change cut-offs. (e) Database for Annotation, Visualization and Integrated Discovery (DAVID) was used to determine the most enriched cellular compartments associated with proteins that were significantly enriched in FBS HI-after EV-depl compared to FBS no-HI. The 10 most enriched terms (based on p-value) are displayed (253 proteins, p-value = 0.001, q-value = 0.009).

The GO term analysis showed that the proteins enriched in the sEV pellet from HI-after EV-depl FBS compared to no-HI FBS were mostly associated with the cellular compartment term “extracellular exosome” followed by “cytosol” and “membrane”. Interestingly, among the top 20 up-regulated proteins in sEVs from HI-after EV-depl FBS versus no-HI FBS (Fig. S7e) and HI-before EV-depl FBS (Fig. S7f) we found proteins involved in endosome maturation, such as vacuolar protein sorting-associated protein 11 (VPS11)^44, 45^ and sorting nexin 6 (SNX6), which has previously been identified in EVs isolated from colorectal cancer cells^46^. To further investigate the enriched proteins in FBS-derived sEVs from HI-after EV-depl FBS, we compared the proteins enriched in sEVs from HI-after EV-depl FBS versus no-HI FBS as well as HI-after EV-depl FBS versus HI-before EV-depl FBS to proteins identified in the online EV database Vesiclepedia^47^ (Fig. 7). Interestingly, 30 proteins were in common between all three groups, and among them 8 proteins were found in the top 100 proteins of Vesiclepedia (CCT5, EIF4A1, IQGAP1, PDCD6IP, PKM, RAC1, YWHAB, and YWHAG).

**FIGURE 7.**
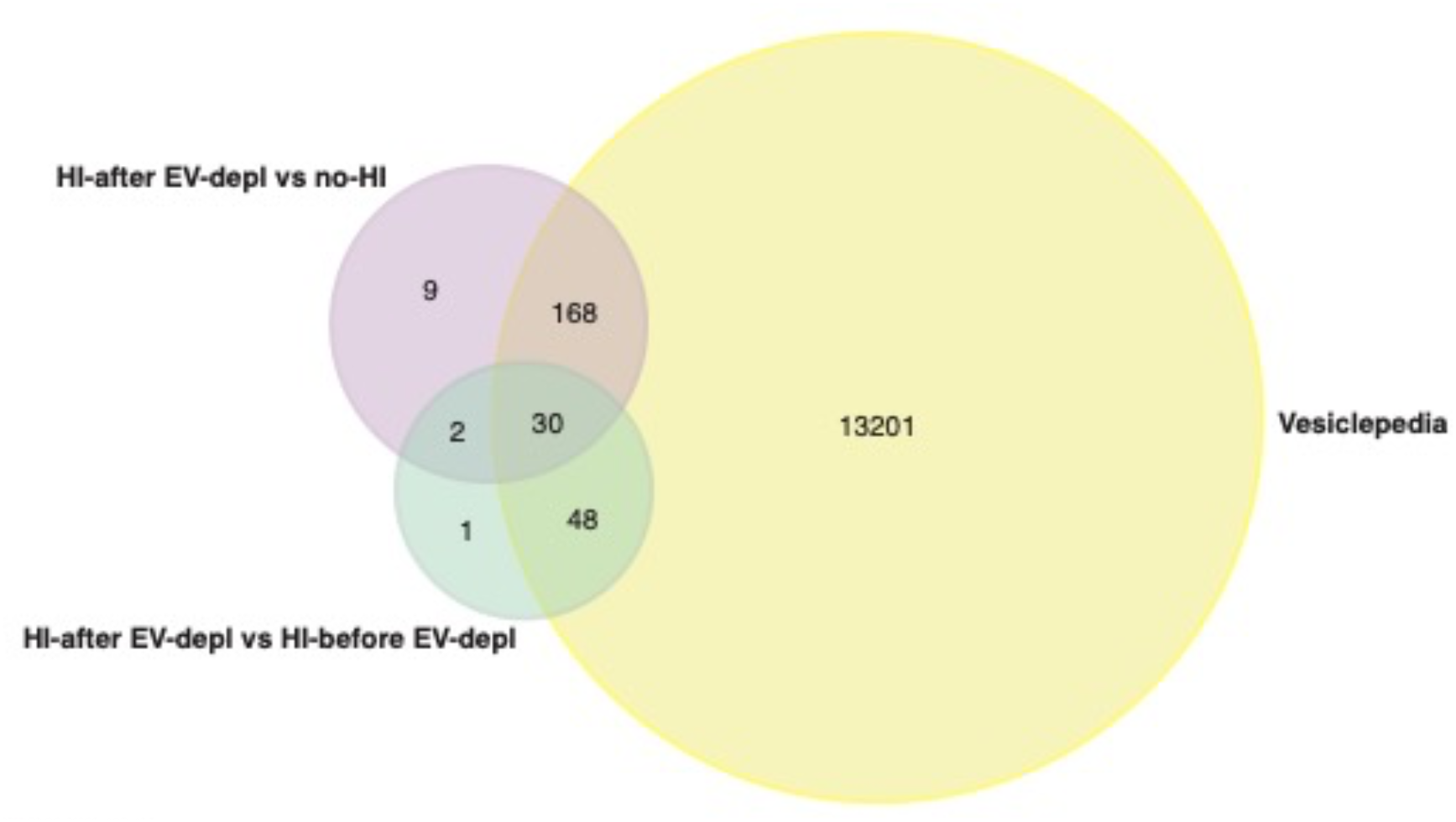
Comparison of proteins up-regulated in sEVs isolated from HI-after EV-depl FBS versus Vesiclepedia^47^. The Venn diagram was created with Fun Rich.

Moreover, although not listed in the top 100 proteins of Vesiclepedia, other shared proteins were previously found in cell-derived EVs, for example, signal transducer and activator of transcription 3 (STAT-3) and matrix metalloproteinase-19 (MMP-19). STAT-3 is a member of the STAT family and is a signal transducer recently identified in EVs isolated from colorectal cancer cells^48^, and MMP-19 was found in the secretome of mammalian kidney cells expressing YBX1/YB-1^49^, a gene involved in cell proliferation and invasion^50^. Among the 168 EV proteins shared between HI-before EV-depl FBS versus no-HI FBS and Vesiclepedia, we also identified heat shock protein family A member 4 (HSPA4), which is a member of the heat shock protein 70 family and is a known marker of EVs^51^. Interestingly, only 12 proteins were not listed in Vesiclepedia (listed in Supplementary Table 2). Among those proteins, 5’-nucleotidase cytosolic IIIA (NT5C3A) and 5’-nucleotidase cytosolic IIIB (NT5C3B) have been identified in EVs released by reticulocytes^52^ and medulloblastoma cells^53^, respectively. These results suggest that HI performed after EV depletion by UC induced the formation of proteins that ended up in the sEV pellet and may alter the proteomic analysis of cell-derived sEVs.

## 4 DISCUSSION

In this work, we demonstrated that heat inactivation of FBS (56°C for 30 min under agitation) prior to adding it to cell culture media impacts the purity of EVs isolated from cell cultures. We isolated lEVs and sEVs from three cell lines cultured in media supplemented with FBS that underwent no-HI, HI-before EV-depl, and HI-after EV-depl, and we demonstrated that heat inactivated FBS after EV depletion by UC contaminated sEV samples but not lEV samples. Through proteomic analysis, we identified protein contaminants and found that some of the identified proteins could be mistakenly attributed to human cell-derived sEVs.

As reported in our recent work^54^, although FBS is still the most common supplement for cell culture media, its use may become a disadvantage due to the presence of contaminants (endotoxins, viruses, and mycoplasma)^20^ and EVs that can end up in cell-derived EV samples^55^. The most commonly used technique for FBS EV depletion (centrifugation at 100,000 × *g* for 18 h)^17^ has previously been shown to be unable to completely eliminate EVs from FBS^28, 32^. Despite the efforts of the scientific community to standardize the protocols to reduce FBS-derived contaminants in cell-derived EV samples^28^, one aspect that has not be taken in consideration is the heat inactivation procedure. Although some studies have questioned the heat inactivation procedure^56^, this practice remains commonly used in many laboratories. We analysed the top 100 publications in Google Scholar searching for “exosomes OR extracellular vesicles AND FBS AND heat inactivation” in which EVs were isolated from cell culture media. Among the 83 publications in which the authors stated to have performed heat inactivation of FBS, in 24 works the cell media was changed with serum-free media or media supplemented with commercial Exo-free FBS before EV isolation differently from the remaining 59 publications in which heat-inactivated and EV depleted FBS was added to the cell culture used for EV isolation. Among them, in only 5 publications did the authors specify at which step of the FBS EV-depletion protocol they performed the heat inactivation. This highlights that in most publications the authors do not specify whether they heat inactivate FBS before or after EV depletion.

Because some studies have demonstrated that heat inactivation of FBS can impair cell growth and phenotype^57–59^, as a first step we analysed the cell viability and morphology of melanoma cells maintained in media supplemented with FBS obtained following the three different protocols. The reason we performed the experiments on different cell lines was to extend the results obtained to multiple *in vitro* models. While MML-1 is a commercially available melanoma cell line, UM22Bap1^+/+^ and UM22Bap1^-/-^ cells were obtained from a patient with uveal melanoma and were transfected using a retroviral vector with a functional copy of *BAP1* (UM22Bap1^+/+^) or an empty vector control sample (UM22Bap1^-/-^) ^39^. While we did not observe any significant changes in cell viability, the growth of MML-1 and UM22Bap1^-/-^ cells in media with HI-before EV-depl FBS and HI-after EV-depl FBS was slightly reduced compared to media plus no-HI FBS. Previous studies demonstrated that heat-inactivated FBS can affect cell metabolism^58, 59^, and this could be the reason why MML-1 and UM22Bap1^-/-^ cell growth was reduced in media supplemented with HI-before EV-depl FBS and HI-after EV-depl FBS.

Because we observed an increase in the protein concentration of sEV samples obtained from HI-after EV-depl FBS, we performed quantitative mass spectrometry analysis to identify the EV proteins that were up-regulated in sEVs from HI-after EV-depl FBS versus HI-before EV-depl FBS and no-HI FBS. To our knowledge, this is the first study that has investigated the protein content of FBS-derived EVs because other works have focused mainly on complete FBS and its effects on the secretome^60–63^. We mapped the identified peptides against the bovine database – based on the bovine origin of FBS – and against the human database in order to determine whether some FBS-derived EV proteins could be misclassified as human proteins and consequently affect EV proteomic analysis in human cell cultures. We found that both proteins assigned as bovine and human that were isolated from sEVs derived from HI-after EV-depl FBS differed significantly from sEVs derived from no-HI FBS and HI-after EV-depl FBS. Although there are no studies in the literature that have directly compared the protein contents of FBS before and after heat inactivation, it is known that the proteome of cells cultured in media supplemented with heat-inactivated FBS can differ from the same cells grown in media supplemented with non-heat-inactivated FBS ^64, 65^, thus suggesting that the FBS protein content is affected by the heat inactivation procedure. Interestingly, it has recently been demonstrated that FBS heat inactivation causes the denaturation of the protein corona due to structural changes induced by the increased temperature, thus reducing the binding affinity of the serum proteins^66^. This may explain the differences that we observed in our three conditions (no-HI, HI-before EV-depl, and HI-after EV-depl) because if some proteins are bound or unbound to each other they might precipitate and pellet differently during the UC process.

The search for FBS-derived EV proteins in the human database identified several proteins that could be attributed to human origin. This result could be explained by the existing homology between the human and bovine species^67^. Previous studies demonstrated that human antibodies can incorrectly recognize bovine proteins. In particular, Shelke et al. detected the presence of TSG101, CD81, and CD63 in EVs isolated from FBS using human antibodies, thus highlighting the very high homology between humans and bovines^28^. Importantly, most of the proteins that were up-regulated in FBS-derived EVs from HI-after EV-depl FBS versus no-HI FBS and HI-before EV-depl FBS have been previously described in cell-derived EV proteomics studies^49, 51^. These could at least partly be due to FBS and mistakenly attributed to human cell line-derived EVs. Eight of these proteins were found in the top 100 proteins in Vesiclepedia^47^, which is the most commonly used database for studies on EV proteins. Among these eight proteins, the mRNA helicase EIF4A1 has been identified in EVs isolated from several cell lines, including adipocytes^68^, breast cancer cells^69^, and fibroblasts^70^. Moreover, a type of metalloproteinase – MMP-19 – was found among the proteins that were up-regulated in FBS-derived sEVs from HI-after EV-depl FBS. Matrix metalloproteinases are known to be enriched in EVs^71^, and in particular the presence of MMP-1, −14, and −19 was previously observed in EVs derived from Madin–Darby canine kidney cells^72^.

Of note, the different outputs between bovine and human analyses could be due to the lack of studies regarding bovine proteins, making the bovine datasets less representative compared to human datasets. This might explain why we did not find the category “extracellular exosomes” in the DAVID analysis of bovine proteins because the GO term “extracellular exosome” has over 3,000 proteins assigned to it in the human database but only 74 proteins in the bovine database.

Our results are strictly focused on the methodologies and proteomic analyses. It would be of great interest to investigate the functional effects of FBS-derived sEVs from HI-after EV-depl FBS in comparison to FBS-derived sEVs from no-HI FBS and HI-before EV-depl FBS to determine if the effects are due to FBS-derived sEVs or to protein contaminants induced by heat inactivation that end up in the EV pellet.

## 5 CONCLUSION

In this study we highlighted for the first time the contribution of FBS in EV samples linked to the timing of FBS heat inactivation. We demonstrated that heat inactivation of FBS can reduce EV purity, increase the amount of EV-derived proteins in cell cultures, and confound subsequent proteomic analysis. We recommend that EV researchers specify whether they perform heat inactivation of FBS in their studies, and we strongly recommend to either not perform heat inactivation of FBS or, if it is necessary, to perform heat inactivation prior to EV depletion.

## AUTHOR COTRIBUTION

RC designed the study, performed the TEM experiments, analysed the data and provided funding. MM and OU performed the EV isolation from cell lines, protein quantification, and particle measurement experiments. CL performed the proteomics data analysis and gave conceptual advice. JJ performed the flow cytometry experiments and flow cytometry analysis. DD helped with EV isolation from FBS and with experiments on cell viability. ROB gave conceptual advice and provided funding. The manuscript was written by OU and RC with the support of CL and ROB. All authors have read, commented on, and given approval of the final version of the manuscript.

## ACKNOWLEDGEMENTS

We acknowledge the Centre for Cellular Imaging at the University of Gothenburg and the National Microscopy Infrastructure (VR-RFI 2016-00968) for providing assistance in TEM. For proteomic analysis we thank Annika Thorsell at the Proteomics Core Facility at Sahlgrenska Academy, University of Gothenburg. We thank Jonas Nilsson and Lisa Nilsson for kindly sharing the UM22Bap1^+/+^ and UM22Bap1^-/-^ cell lines. The authors acknowledge Roberto Cattaneo for his assistance with graphic design.

## FIGURES

**SUPPLEMENTARY FIGURE 1.**
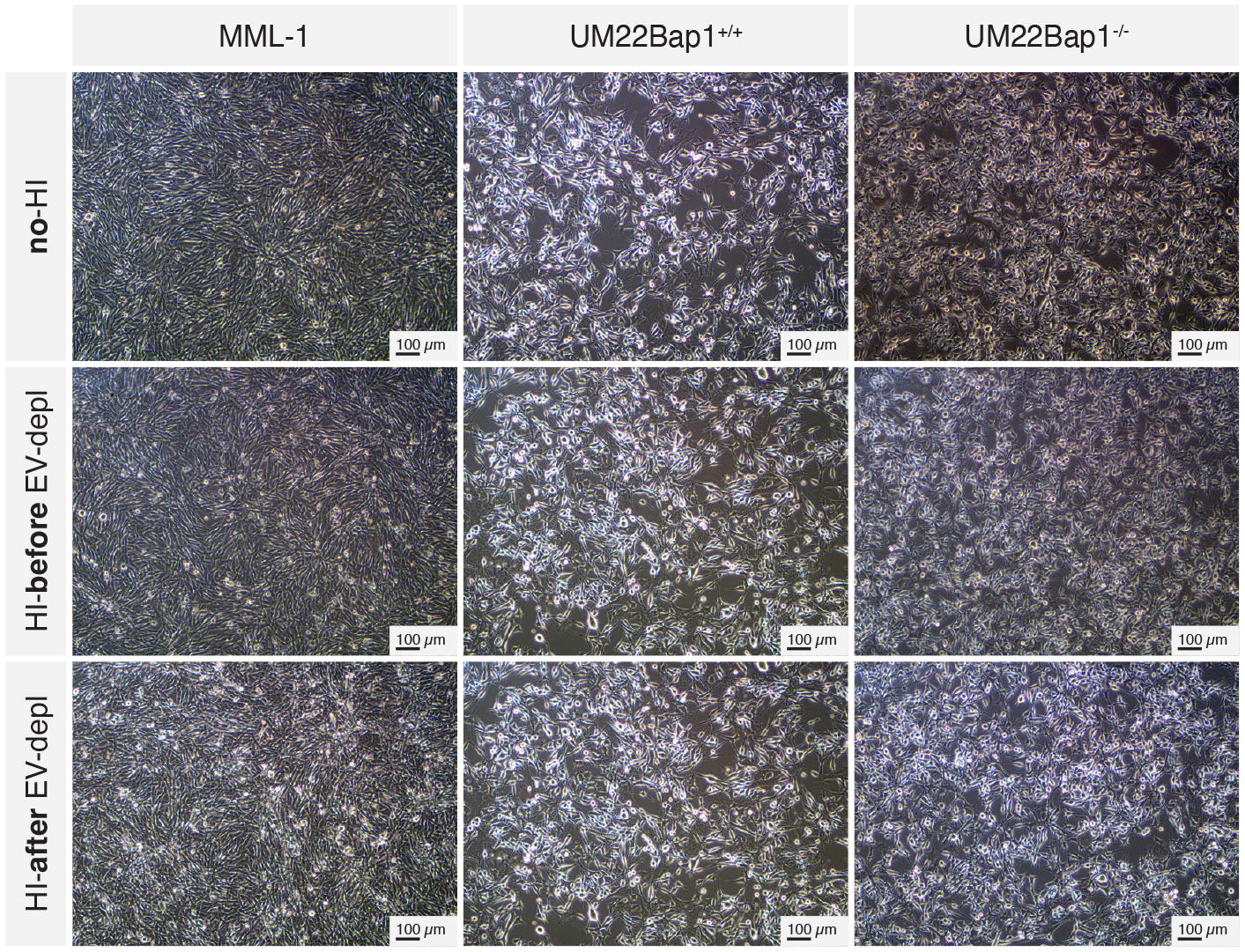
Visualization of cell confluence. Light microscope images of MML-1, UM22Bap1^+/+^, and UM22Bap1^-/-^ cells in media supplemented with no-HI FBS, HI-before EV-depl FBS, and HI-after EV-depl FBS. Representative images of three experiments are shown.

**SUPPLEMENTARY FIGURE 2.**
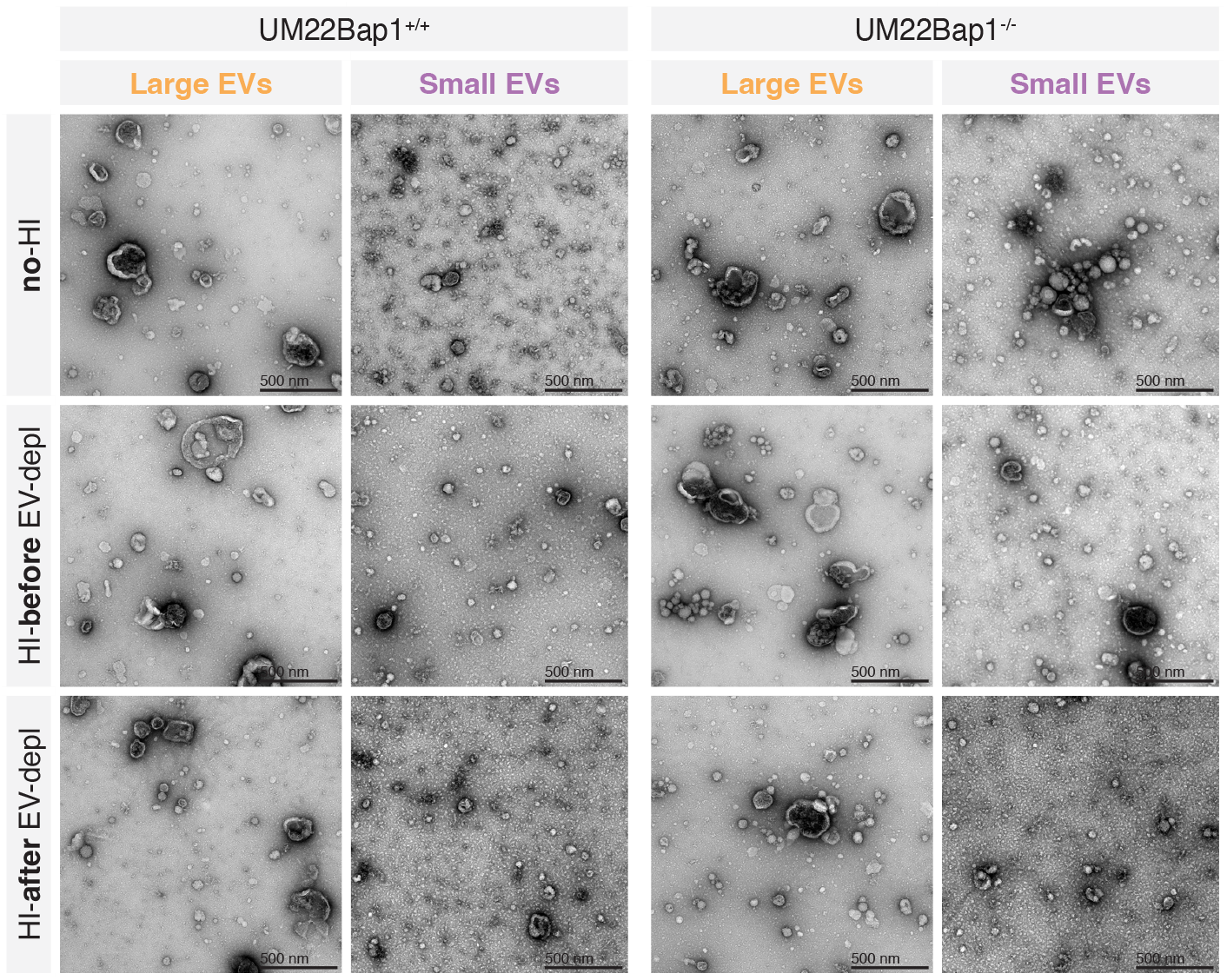
TEM analysis of EVs isolated from UM22Bap1^+/+^ (left panel) and UM22Bap1^-/-^ (right panel) cells cultured in media supplemented with no-HI FBS, HI-before EV-depl FBS, and HI-after EV-depl FBS. A total of 5 μg of lEVs and sEVs were loaded onto grids, negative stained, and evaluated with TEM. Scale bars are 500 nm.

**SUPPLEMENTARY FIGURE 3.**
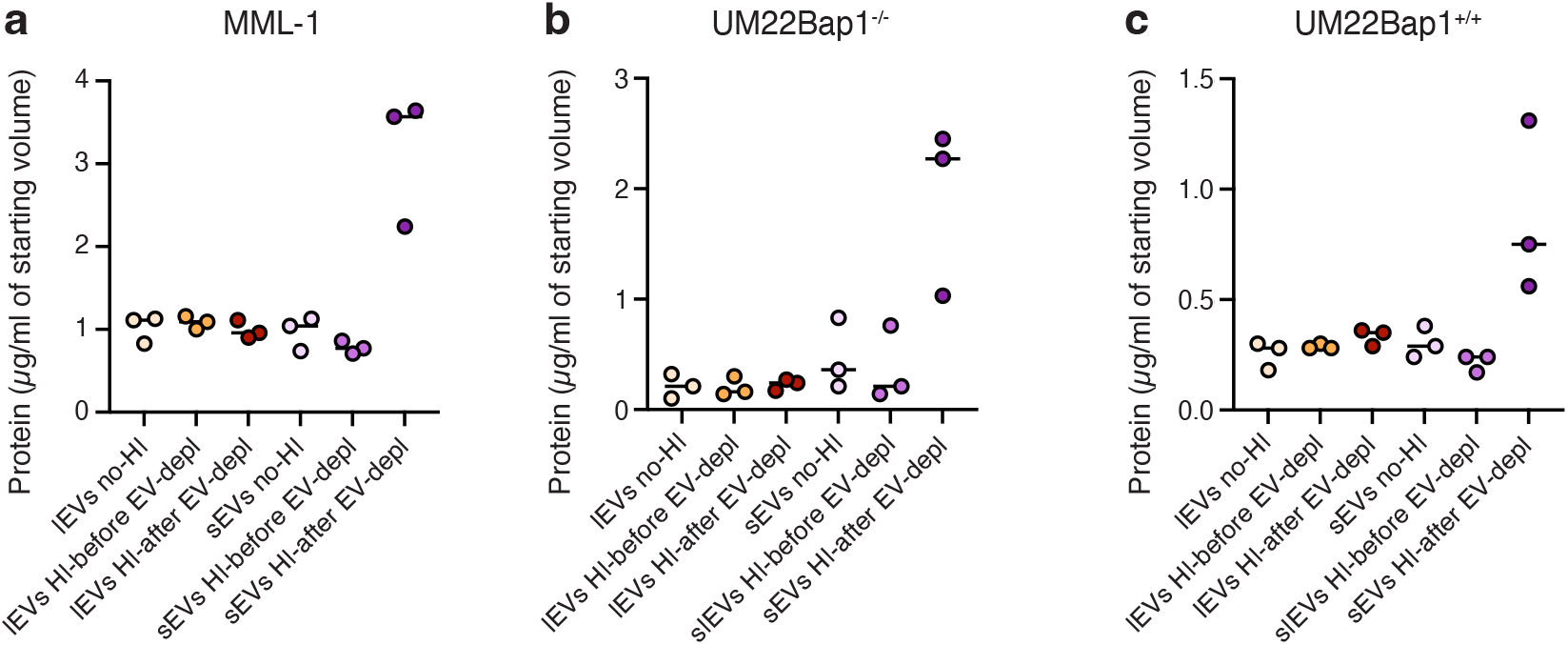
Analysis of EVs isolated from MML-1, UM22Bap1^+/+^, and UM22Bap1^-/-^ cells cultured in media supplemented with no-HI FBS, HI-before EV-depl FBS, and HI-after EV-depl FBS. Protein concentrations of lEVs and sEVs isolated from (a) MML-1, (b) UM22Bap1^+/+^, and (c) UM22Bap1^-/-^ cells in media supplemented with no-HI FBS, HI-before EV-depl FBS, and HI-after EV-depl FBS. The results are presented as the mean, but individual values from three different experiments are shown as well. *N* = 3.

**SUPPLEMENTARY FIGURE 4.**
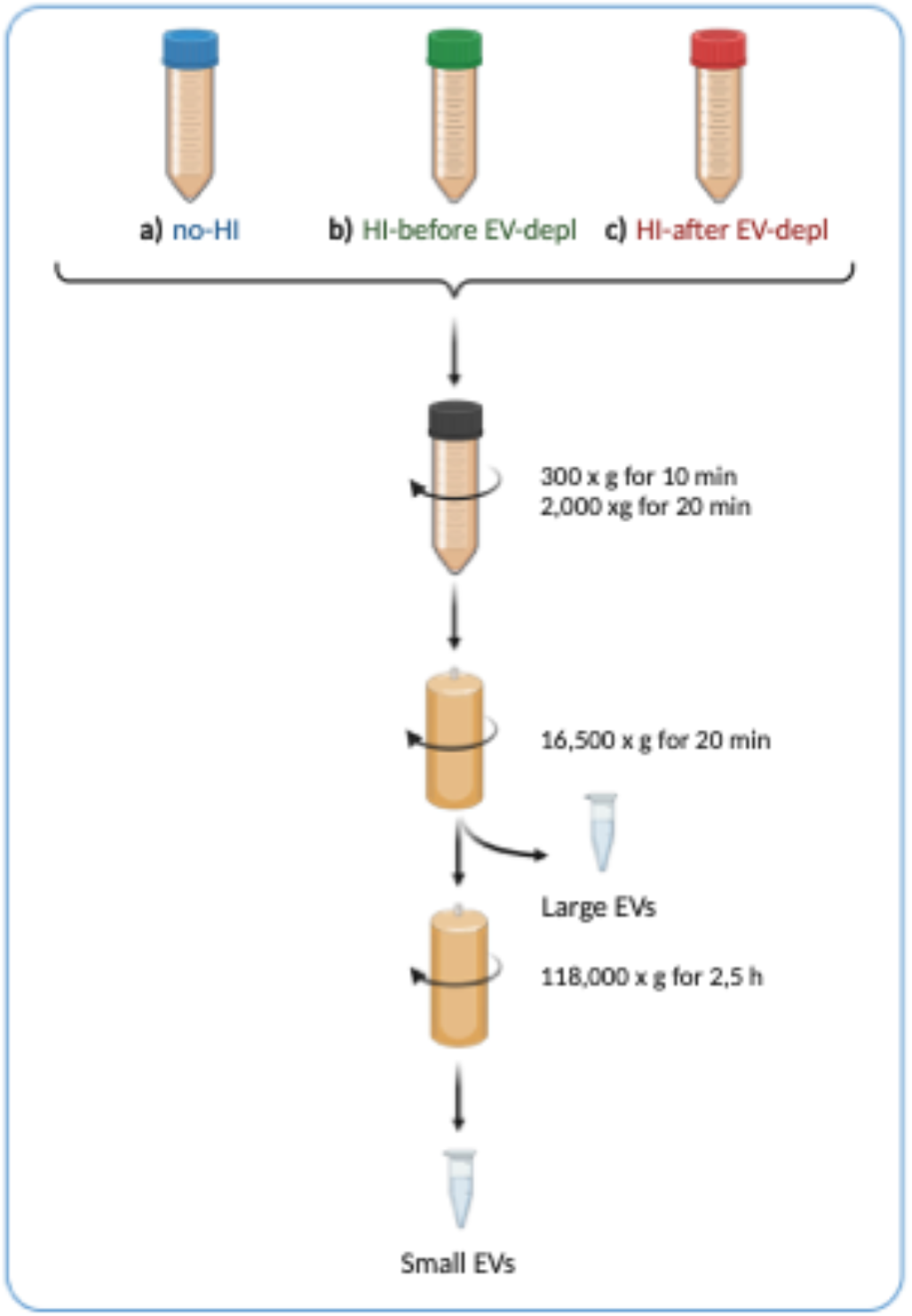
Isolation of lEVs and sEVs from no-HI FBS, HI-before EV-depl FBS, and HI-after EV-depl FBS. Created with BioRender.com

**SUPPLEMENTARY FIGURE 5.**
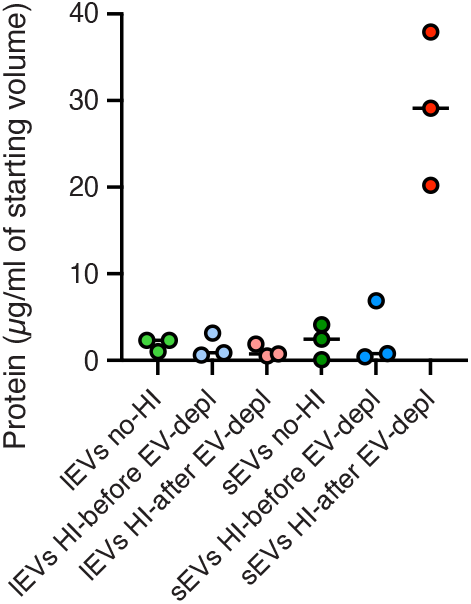
Protein concentration of lEVs and sEVs isolated from no-HI FBS, HI-before EV-depl FBS, and HI-after EV-depl FBS. The results are presented as the mean, but individual values from three different experiments are shown as well. *N* = 3.

**SUPPLEMENTARY FIGURE 6.**
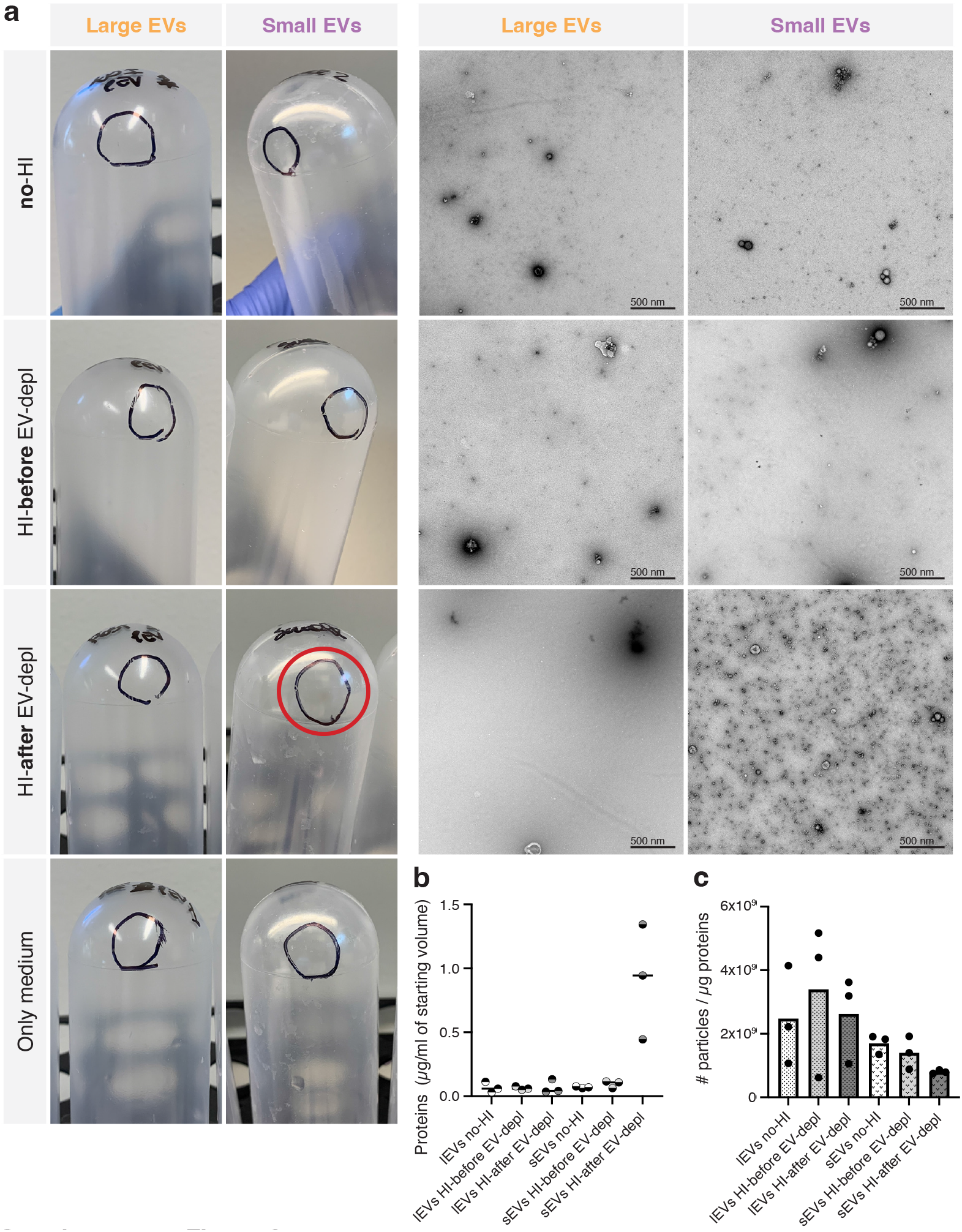
Analysis of pellets obtained by the differential centrifugation of media supplemented with no-HI FBS, HI-before EV-depl FBS, and HI-after EV-depl FBS. (a, left panel) Representative images of pellets obtained by the UC of media supplemented with no-HI FBS, HI-before EV-depl FBS, and HI-after EV-depl FBS. (a, right panel) TEM images of lEVs and sEVs isolated from media supplemented with no-HI FBS, HI-before EV-depl FBS, and HI-after EV-depl FBS. Five micrograms of lEVs and sEVs were loaded onto grids, negative stained, and evaluated with TEM. Scale bars are 500 nm. (b) Protein concentration of lEVs and sEVs isolated from media supplemented with no-HI FBS, HI-before EV-depl FBS, and HI-after EV-depl FBS. (c) The ratio of the particle number and protein concentration of lEVs and sEVs isolated from media supplemented with no-HI FBS, HI-before EV-depl FBS, and HI-after EV-depl FBS. The results are presented as the mean, but individual values from three different experiments are shown as well. *N* = 3.

**SUPPLEMENTARY FIGURE 7.**
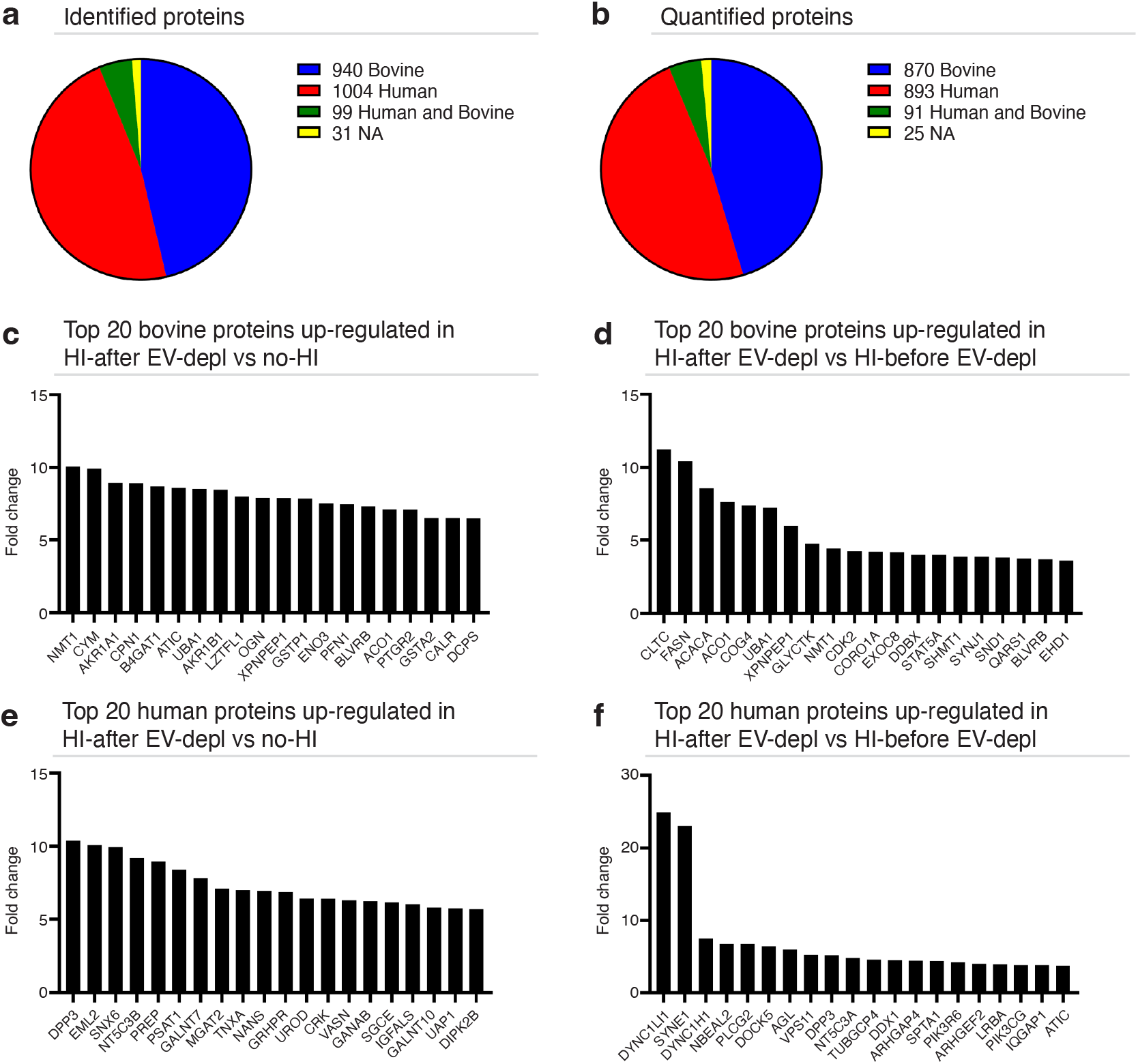
Proteomic analysis of sEVs isolated from no-HI FBS, HI-before EV-depl FBS, and HI-after EV-depl FBS. Pie charts according to species of (a) identified and (b) quantified proteins in sEVs through mass spectrometry (TMT) analysis. (c) The top 20 proteins quantified against the bovine list and up-regulated in sEVs isolated from HI-after EV-depl FBS versus sEVs from no-HI FBS. (d) The top 20 proteins quantified against the bovine list and up-regulated in sEVs isolated from HI-after EV-depl FBS versus HI-before EV-depl FBS. (e) The top 20 proteins quantified against the human list and up-regulated in sEVs isolated from HI-after EV-depl FBS versus sEVs from no-HI FBS. (f) The top 20 proteins quantified against the human list and up-regulated in sEVs isolated from HI-after EV-depl FBS versus HI-before EV-depl FBS.

**TABLE S1.** List of proteins found in both the bovine and human lists.

| Proteins present in both Bovine and Human datasets |  |
| --- | --- |
| 1 | 4-trimethylaminobutyraldehyde dehydrogenase |
| 2 | 6-phosphogluconolactonase |
| 3 | Albumin |
| 4 | Alpha-2-macroglobulin |
| 5 | Alpha-fetoprotein |
| 6 | Annexin A4 |
| 7 | Annexin A5 |
| 8 | Argininosuccinate synthase |
| 9 | Bifunctional purine biosynthesis protein |
| 10 | Cadherin-2 |
| 11 | Cartilage oligomeric matrix protein |
| 12 | Catalase |
| 13 | Coagulation factor V |
| 14 | Coatomer subunit alpha |
| 15 | Collagen alpha-1(II) chain |
| 16 | Complement C3 |
| 17 | Divergent protein kinase domain 2 |
| 18 | Glyceraldehyde-3-phosphate dehydrogenase |
| 19 | Hemoglobin subunit alpha |
| 20 | Hemoglobin subunit beta |
| 21 | Hepatocyte growth factor-like protein |
| 22 | Inter-alpha-trypsin inhibitor heavy chain H3 |
| 23 | Lactotransferrin |
| 24 | Mimecan |
| 25 | Neural cell adhesion molecule 1 |
| 26 | Nicotinate phosphoribosyltransferase |
| 27 | Peroxiredoxin-6 |
| 28 | Prolyl endopeptidase FAP |
| 29 | Protein S100-A9 |
| 30 | Retinol-binding protein 4 |
| 31 | Suprabasin |
| 32 | Thrombospondin-1 |
| 33 | Transketolase |
| 34 | Tubulin beta-6 chain |
| 35 | Ubiquitin carboxyl-terminal hydrolase 15 |
| 36 | Zinc-alpha-2-glycoprotein |

**TABLE S2.**
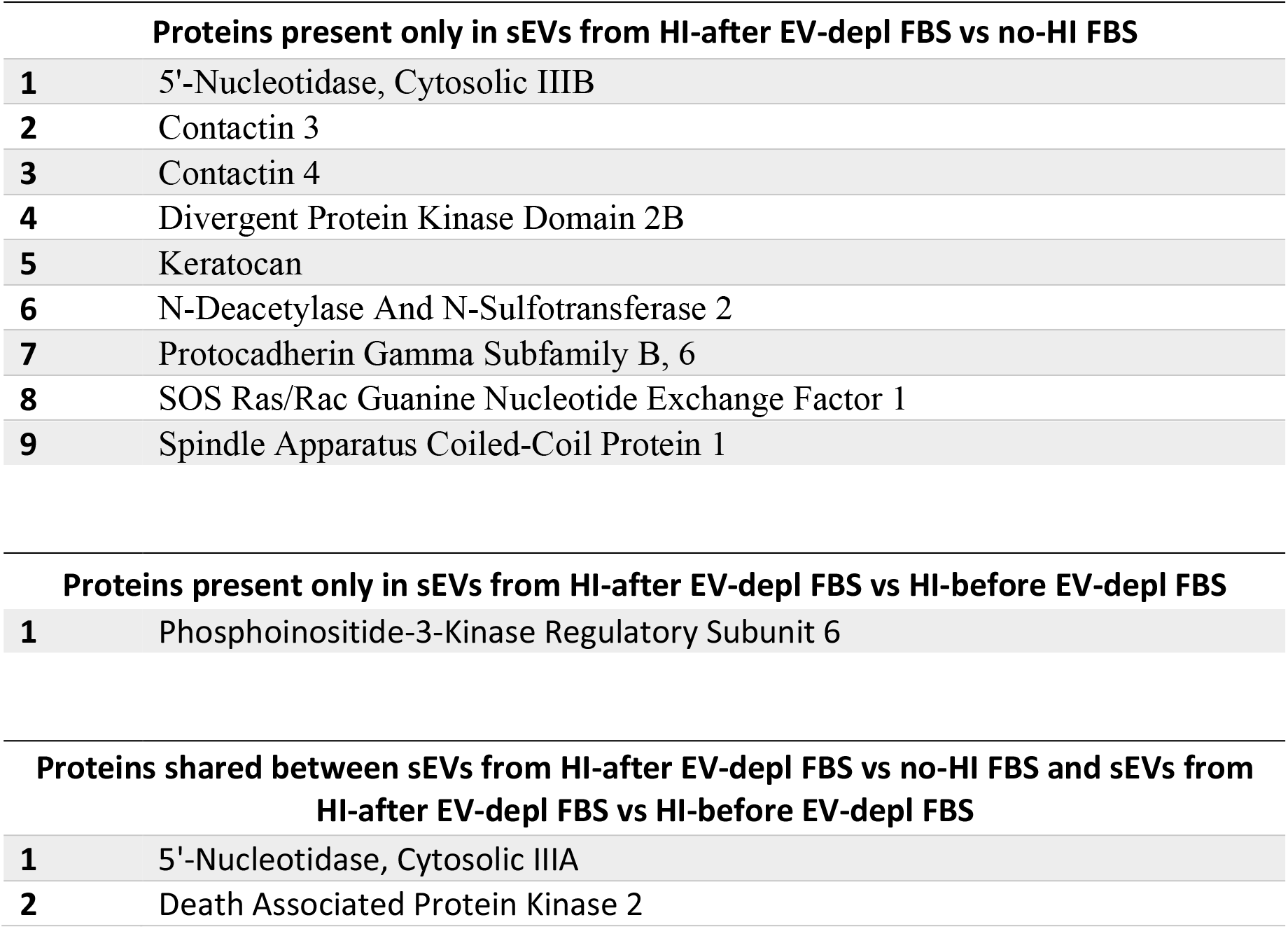
List of human proteins present in FBS-derived sEVs but not in Vesiclepedia^47^.

